# Expression pattern of nicotinic acetylcholine receptor subunit transcripts in neurons and astrocytes in the ventral tegmental area and locus coeruleus

**DOI:** 10.1101/2023.04.28.537363

**Authors:** Runbo Gao, Amelia M. Schneider, Sarah M. Mulloy, Anna M. Lee

## Abstract

Acetylcholine is the endogenous agonist for the neuronal nicotinic acetylcholine receptor (nAChR) system, which is involved in attention, memory, affective behaviors, and substance use disorders. Brain nAChRs are highly diverse with 11 different subunits that can form multiple receptor subtypes, each with distinct receptor and pharmacological properties. Different neuronal cell types can also express different nAChR subtypes, resulting in highly complex cholinergic signaling. Identifying which nAChR subunit transcripts are expressed in cell types can provide an indication of which nAChR combinations are possible and which receptor subtypes may be most pharmacologically relevant to target. In addition to differences in expression across cell types, nAChRs also undergo changes in expression levels from adolescence to adulthood. In this study, we used fluorescent *in situ* hybridization to identify and quantify the expression of α4, α5, α6, β2 and β3 nAChR subunit transcripts in dopaminergic, GABAergic, glutamatergic, and noradrenergic neurons and astrocytes in the ventral tegmental area (VTA) and locus coeruleus (LC) in adult and adolescent, male and female C57BL/6J mice. There were distinct differences in the pattern of nAChR subunit transcript expression between the two brain regions. LC noradrenergic neurons had high prevalence of α6, β2 and β3 expression, with very low expression of α4, suggesting the α6(non-α4)β2β3 receptor as a main subtype in these neurons. VTA astrocytes from adult mice showed greater prevalence of α5, α6, β2 and β3 transcript compared with adolescent mice. These data highlight the complex nAChR expression patterns across brain region and cell type.

## Introduction

Acetylcholine is the endogenous agonist for the neuronal nicotinic acetylcholine receptor (nAChR) system. The nAChRs are widely expressed in the brain and are involved in drug reward responses, attention, memory, affective behaviors, pain processing and inflammation (Picciotto et al., 2008; De Biasi and Dani, 2011; Hurst et al., 2013; Galvin et al., 2020). nAChRs are pentameric ligand-gated cation channels that are primarily located on pre-synaptic terminals and soma, thus these receptors modulate cell excitability and neurotransmitter release. Activation of nAChRs by ACh or by exogenous ligands allows influx of cations (sodium, potassium and calcium) into the cell, leading to a more depolarized state and increased cell excitability (Wills et al., 2022). nAChRs are highly diverse, with 11 different subunits (α2–7, α9– 10, and β2–4) expressed in the mammalian brain that results in multiple receptor subtypes. Each receptor subtype has distinct receptor properties, ligand binding affinities, calcium permeability and deactivation kinetics that are due to the different subunit compositions of the receptors (for comprehensive reviews, see Albuquerque et al., 2009; Wills et al., 2022).

The identification of nAChR subunit composition in the brain has been hampered by the lack of tools to identify individual subunits in cell types and neural circuits. Cells can express multiple nAChR subunits (Azam et al., 2002) and protein detection tools such as Western blotting or radioligand binding lack the specificity required to distinguish subunits. Some nAChR subunits such as the α4 and α7 subunit are widely expressed throughout the brain and are nearly ubiquitous. Less widely expressed subunits, such as α6 and β3, were originally thought to be absent from many brain regions or cell types due to limitations in past detection techniques. An alternative strategy is to probe for nAChR subunit transcripts. Identifying the presence of subunit transcripts can and has been used to provide an indication of which nAChR subtype combinations are possible in specific cell types (Azam et al., 2002; Yan et al., 2018). Technical advances have significantly improved the detection of individual nAChR transcripts, which has led to revised data on the presence of nAChR subunits in cells that were previously thought to be lacking these subunits, such as expression of the α6 subunit in GABA neurons of the ventral tegmental area (VTA) (Charpantier et al., 1998; Steffensen et al., 2018; Wadsworth et al., 2023).

nAChRs containing the α4 and/or the α6 subunits (denoted α4* and/or α6*, with * indicating the presence of other undetermined subunits in the receptor pentamer) have high affinity for nicotine. The α4 subunit is frequently expressed with the β2 subunit in the same pentamer receptor, and the α6 subunit is frequently expressed with the β3 subunit in the same subtype. These α4β2* and α6β3* nAChR subtypes are of interest in substance use disorder (SUD) research, particularly alcohol use disorder (AUD) and nicotine dependence (Grady et al., 2007; Gotti et al., 2010; Hendrickson et al., 2013; Wills et al., 2022). Past studies using global genetic deletions of α4, β2 or α6 subunits individually have demonstrated a role for each subunit in mediating nicotine self-administration, alcohol consumption, and nicotine and alcohol reward (Picciotto et al., 1998; Pons et al., 2008; Hendrickson et al., 2011; Liu et al., 2013; Guildford et al., 2016). Currently, there are no cholinergic drugs that are FDA-approved for the treatment of AUD. Varenicline is approved for nicotine dependence and was designed as a partial agonist targeting the α4β2 subtype (Rollema et al., 2007) but has additional partial agonist activity at the α6* subtype and full agonist activity at the α7 subtype (Bordia et al., 2012; Mihalak et al., 2006). The α4 subunit is widely expressed in rodent and human brain but intriguingly α6 and β3 are heavily expressed in only a small number of brain regions, including the VTA, locus coeruleus (LC), substantia nigra and superior colliculus (Le Novere et al., 1996; Champtiaux et al., 2002; Mackey et al., 2012).

In this study, we focused on the VTA and LC. The VTA is critically important for natural and drug reward processing and is cellularly heterogenous, with dopaminergic, GABAergic, glutamatergic neurons and neurons that co-express two neurotransmitters (Nestler, 2005; Trudeau et al., 2014; Morales and Margolis, 2017). All the brain nAChR subunits have been found in the VTA, excluding α9 and α10. The nAChR subunit composition in VTA dopamine neurons has been previously investigated (Charpantier et al., 1998; Azam et al., 2002; Yang et al., 2011; Ngolab et al., 2015), since VTA dopamine neurons play a critical role in salience and rewarding responses of drugs of abuse (Morel et al., 2019; Hynes et al., 2021). Less known about nAChR subunit composition in VTA GABA and glutamate neurons, and characterization of nAChR subunit expression on different VTA cell types is still lacking. The LC has also been implicated in SUD, AUD, as well as circadian rhythms, aging, and stress responses (Poe et al., 2020; Beardmore et al., 2021; Downs and McElligott, 2022). The LC is primarily considered a noradrenergic region but is also heterogenous and contains GABAergic and glutamatergic neurons. Although the LC has been far less investigated compared with the VTA, rodent studies have detected α3, α4, α6, β2, β3 and β4 nAChR subunits in the LC (Lena et al., 1999; Le Novere et al., 1996; Champtiaux et al., 2002; Mackey et al., 2012). However, little is known about which nAChR subunits are expressed in which LC cell types. In addition to investigating nAChR expression in different neuronal cell types in the VTA and LC, nAChR expression in astrocytes is also of interest. Astrocytes are important regulators of synaptic function and an emerging role for astrocytes in SUD is being recognized (Giacometti et al., 2020; Wang et al., 2022). The expression of nAChR subunit transcripts in astrocytes has been previously reported, with α4 and α7 detected in rat astrocyte cultures (Xiu et al., 2005). However, expression of other nAChR subunits on astrocytes has not been well characterized and little is known about how nAChR subunit expression on astrocytes differs across brain region. In this study, we used fluorescent *in situ* hybridization to determine the expression of individual nAChR subunits in multiple cell types of the VTA and LC and found distinct differences in the pattern of nAChR subunit expression between the two brain regions.

## Methods

### Animals

Male and female C57BL/6J adult and adolescent mice were obtained from The Jackson Laboratory, three per age per sex. Adult mice were 56 days of age and adolescent mice were 21 days of age on arrival. Mice were group housed and acclimated to our facility for 5-7 days prior to euthanization and brain collection. Brains were frozen and stored at -80°C until sectioning.

### *Fluorescent* in situ *hybridization with HiPlex RNAscope*

Coronal sections (16 μM) from the rostral to caudal area of VTA and LC were collected from all mice. Multiple 16 μM sections from each brain region from each mouse were processed and imaged in a nested experimental design. Fluorescent *in situ* hybridization using the HiPlex RNAscope assay (#324108, Advanced Cell Diagnostics, Inc.) was used to detect cell-type-specific expression of nAChR subunit transcripts in the VTA and the LC. The sections were processed using the HiPlex RNAscope protocol according to manufacturer recommendations. In the first step, all target probes were hybridized and amplified simultaneously and remain hybridized with target transcripts throughout the three assay rounds. In each assay round, three different transcript species are revealed with different fluorophores, which are then cleaved off to reveal another three transcript species. The target transcript species were *Chrna4* (Mm-Chrna4 #429871), *Chrna5* (Mm-Chrna5 #312571), *Chrna6* (Mm-Chrna6 #467711), *Chrnb2* (Mm-Chrnb2 #449231), *Chrnb3* (Mm-Chrnb3 #449201), *Slc17a6* (vesicular glutamate transporter 2, Mm-Slc17a6 #319171), *Slc32a1* (GABA vesicular transporter, Mm-Slc32a1 #319191), *Th* (tyrosine hydroxylase, Mm-Th #317621) and *Gfap* (Glial fibrillary acidic protein, Mm-Gfap #313211). 4’,6-diamidino-2-phenylindole (DAPI) stain was included for all slides. Slides were mounted with Prolong Gold Anti-Fade Reagent (#9071, Cell Signaling Technology), cover slipped and stored at 4°C in the dark before imaging.

### Image acquisition and analysis

Fluorescence images were captured at 40X using a Keyence microscope (BZ-X700 series). Each region of interest (ROI) consisted of a left or right section from VTA or LC, with ROIs from 3 different mice per sex per age. Each image was loaded into MATLAB Computer Vision Toolbox (MathWorks; Natick, MA) for image registration followed by ROI selection. The VTA and LC images were stitched as necessary to encompass the entire ROI region. Locations of VTA and LC were recorded so that the same field of view could be located in all three rounds of imaging in the assay. The ROIs for all the VTA images consisted of an identically sized rectangle across all sections and animals, capturing the medial VTA within the area of positive tyrosine hydroxylase transcript (*Th*) expression, a marker for dopamine neuron expression in the VTA. ROIs for the LC were traced outlines of the LC based on *Th* transcript expression, a marker for noradrenergic neuron expression in the LC. Registered ROI image sets were exported from MATLAB to CellProfiler (www.cellprofiler.org, RRID:SCR_007358) (Carpenter et al., 2006; Kamentsky et al., 2011; McQuin et al., 2018; Stirling et al., 2021). The CellProfiler program was first used to subtract background fluorescence, and intensity information was split into HSV color space and converted to grayscale based on the intensity of each pixel, and the cell areas were automatically identified in an unbiased manner based on DAPI expression and the Ostu thresholding method. CellProfiler program was then used to relate the signal punctum to cell area for each probe in an unbiased manner to generate the number of probe puncta per cell in each ROI. Data were then imported into MATLAB, where the background probe rate for each probe-fluorophore combination was determined using puncta expression in the non-cell area. The probability that a cell was positive for a target probe was calculated using a binomial distribution test comparing the background probe expression and the puncta expression within the cell area, with an α of 0.01 and a Bonferroni correction using the number of cells in that ROI. The percentage of cells per ROI that were positive for a single probe or positive for two probes of interest was then calculated. ROIs with fewer than 75 cells were excluded from the analyses.

### Statistical analysis

All statistical analyses were performed with Prism 9.5.1 (GraphPad Software). Outliers were identified as ROIs with percentage cell or transcript expression that was outside twice the standard deviation from the mean, or identified with the Grubbs test, and were omitted from the analyses. For the VTA data, we omitted one ROI from one adolescent male and one ROI from 1 adult male. For the LC data, we omitted one ROI from one adolescent male, one ROI from one adult male, one ROI from one adult female and one ROI from one adolescent female. We then confirmed that the data were normally distributed, allowing for parametric analyses. As the data were obtained with a hierarchical sampling design with multiple ROIs imaged from each animal, we used hierarchical (nested) analyses (a nested two-tail Student’s *t*-tests to compare 2 groups or a nested one-way ANOVA to compare 2+ groups) to account for non-independence of the data points. To determine whether the number of cells per ROI differed across all four groups in the VTA and LC, a nested ANOVA was used and data were expressed as mean ± standard deviation (SD). All transcript expression data (cell-type markers and nAChR subunits) are presented as the percentage of cells per ROI expressing the target transcript. We investigated the effects of age in males and females separately. If there were no effects of age, adults and adolescents were combined to examine differences between males and females for a subset of comparisons. All transcript data are expressed as mean ± standard error (SEM).

## Results

### Single and co-expressing cell types in the VTA

In this study, we used an unbiased multiplex fluorescent *in situ* hybridization approach (HiPlex RNAscope) to identify the expression of α4, α5, α6, β2 and β3 nAChR subunit transcripts in dopaminergic, GABAergic, glutamatergic and noradrenergic neurons and in astrocytes of the VTA and LC in adult and adolescent, male and female C57BL/6J mice. A schematic of the regions of interest (ROI) and representative images from one assay round of the VTA and LC are shown in **Fig. 1**.

**Fig. 1.**
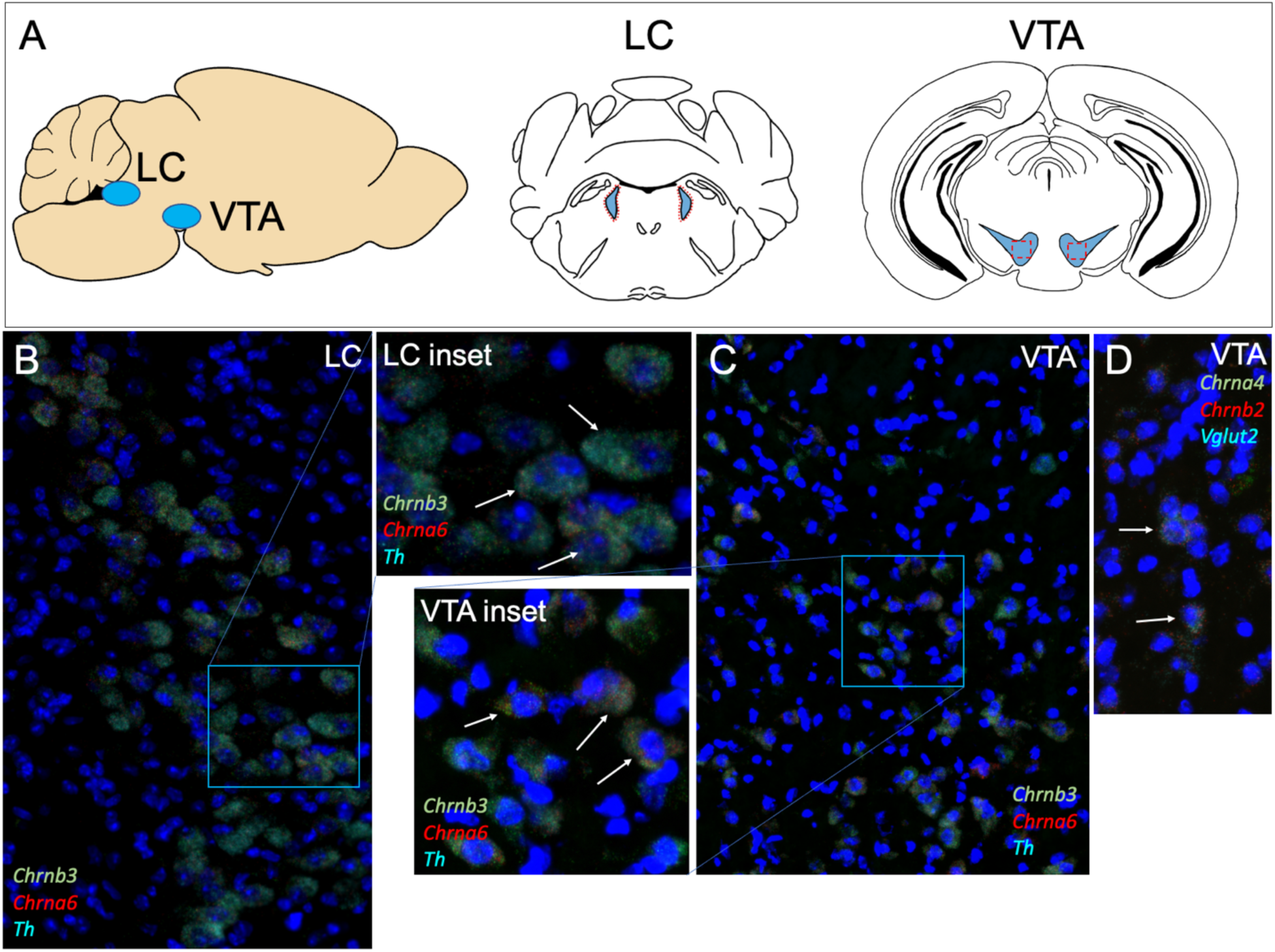
**(A)** *Left:* Schematic of the ventral tegmental area (VTA) and locus coeruleus (LC) in the mouse brain. *Middle, right:* ROIs for each brain region are outlined in red dotted lines. **(B-D)** Representative *in situ* hybridization images obtained after one assay round using fluorescent *in situ* hybridization with HiPlex RNAscope in the LC and VTA. DAPI stain was used to identify cell nuclei in blue. Probes for **B,C**: β3 transcript *(Chrnb3)* in green, α6 transcript *(Chrna6)* in red and tyrosine hydroxylase transcript *(Th)* in cyan. Probes for **D**: α4 transcript *(Chrna4)* in green, β2 transcript *(Chrnb2)* in red and vesicular glutamate transporter *(Vglut2)* in cyan. Arrows point to examples of cells that have positive expression of three fluorescent probes.

We first determined whether the ROIs from the VTA contained equal number of cells across all groups. The average number of cells per VTA ROI was 489.4 ± 142.5 (mean ± SD) with no significant difference between adolescent males, adult males, adolescent females and adult females in the VTA (nested one-way ANOVA, F=0.56, *P*=0.65, *n*=4-8 ROIs per mouse, *n*=3 mice per sex per age). We then examined cell type expression in the VTA starting with dopaminergic cells. The percentage of cells per ROI positive for *Th* (%*Th*+), a marker of dopaminergic cells in the VTA, was similar among all groups with 29.4 ± 0.8% in adolescent males, 30.9 ± 1.1% in adult males, 28.0 ± 1.0% in adolescent females, and 29.9 ± 1.2% in adult females (mean ± SEM, **Fig. 2A**). The percentage of cells positive for *Slc32a1* (%*Vgat+*) transcript, a GABA cell type marker, was similar among all groups with 18.1 ± 0.6% in adolescent males, 19.2 ± 0.8% in adult males, 21.0 ± 0.7% in adolescent females and 19.2 ± 0.9% in adult females (**Fig. 2B**). There were no significant differences between age for either sex or between sex when age was collapsed for either *Th*+ or *Vgat*+ cells. The percentage of cells positive for *Slc17a6* (%*Vglut2*+), a glutamatergic cell type marker, was 5.9 ± 0.5% in adolescent males, 7.0 ± 0.5% in adult males, 7.8 ± 0.4% in adolescent females and 9.3 ± 0.8% in adult females. There were no differences between age within sex. When age was collapsed we found that females had a small but significantly higher %*Vglut2+* cells compared with males (nested Student’s *t*-test, *t*=3.08, \**P*=0.01, **Fig. 2C**). For the percentage of cells positive for *Gfap* (%*Gfap+*), a marker of astrocytes, we found that adult males had a significantly higher percentage of astrocytes compared with adolescent males (*t*=4.60, *P*=0.01), with 11.1 ± 0.6% in adult males and 7.4 ± 0.4% in adolescent males. Females did not show a significant difference in %*Gfap*+ cells across age (9.4 ± 0.5% in adult females and 7.8 ± 0.4% in adolescent females, *P*=0.11, **Fig. 2D**). Overall, these data indicate that male adult mice have a greater proportion of astrocytes in the VTA compared with adolescents, and that age does not influence the expression of neuronal cell types in the VTA.

**Fig. 2.**
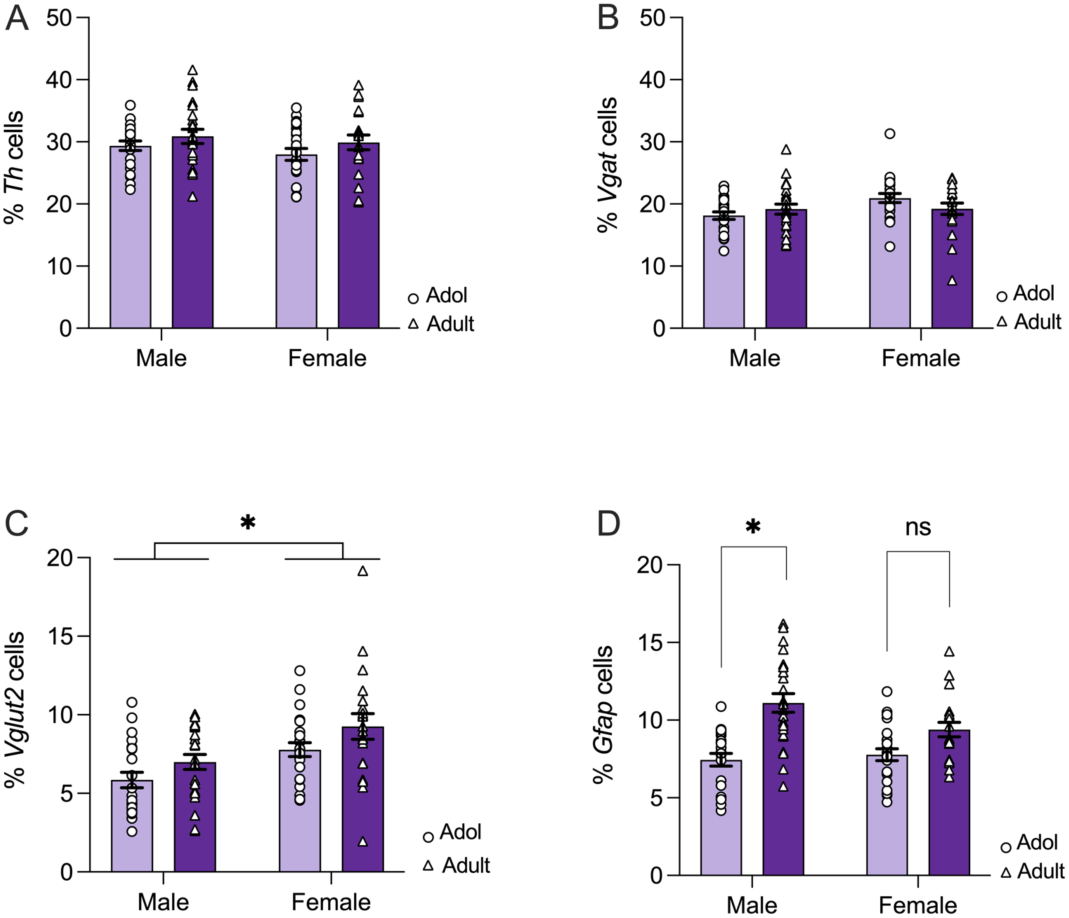
The percentage of cells in VTA ROIs that were positive for **(A)** *Th* and **(B)** *Vgat* transcript were similar across groups. **(C)** Female mice had a higher percentage of *Vglut2* expressing cells compared with males (\**P*=0.01). **(D)** Male adult mice had a higher percentage of *Gfap+* cells compared with male adolescents (\**P*=0.01). There was no significant difference between adult females and adolescent females (ns, *P*=0.11). Data expressed as mean ± SEM, with 4-8 ROI per mouse from *n*=3 mice per sex per age.

We then examined cells that were positive for two neurotransmitters, which indicates potential neurotransmitter co-expression, in the VTA. There were no significant differences in %*Th*+*Vgat*+ or %*Th*+*Vglut2*+ prevalence between ages in either males or females or between sex when age was collapsed (all *P*>0.05). The percentage of %*Th*+*Vgat*+ was 5.6 ± 0.7% in adolescent males, 4.8 ± 0.2% in adult males, 5.5 ± 0.5% in adolescent females and 5.6 ± 0.5% in adult females (**Fig. 3A**). The %*Th*+*Vglut2*+ was 3.7 ± 0.4% in adult males, 2.2 ± 0.2% in adolescent males, 4.8 ± 0.6% in adult females and 3.3 ± 0.3% in adolescent females (**Fig. 3B**). For %*Vgat+Vglut2+* (**Fig. 3C**) we found no differences between age in either males or females (1.6 ± 0.2% in adult males, 1.4 ± 0.2% in adolescent males, 2.5 ± 0.4% in adult females and 2.6 ± 0.3% in adolescent females). When age was collapsed, we found a small but significant difference between sex, with females having a higher %*Vgat+Vglut2+* cells compared with males (nested Student’s *t*-test, *t*=3.02*, P*=0.01).

**Fig. 3.**
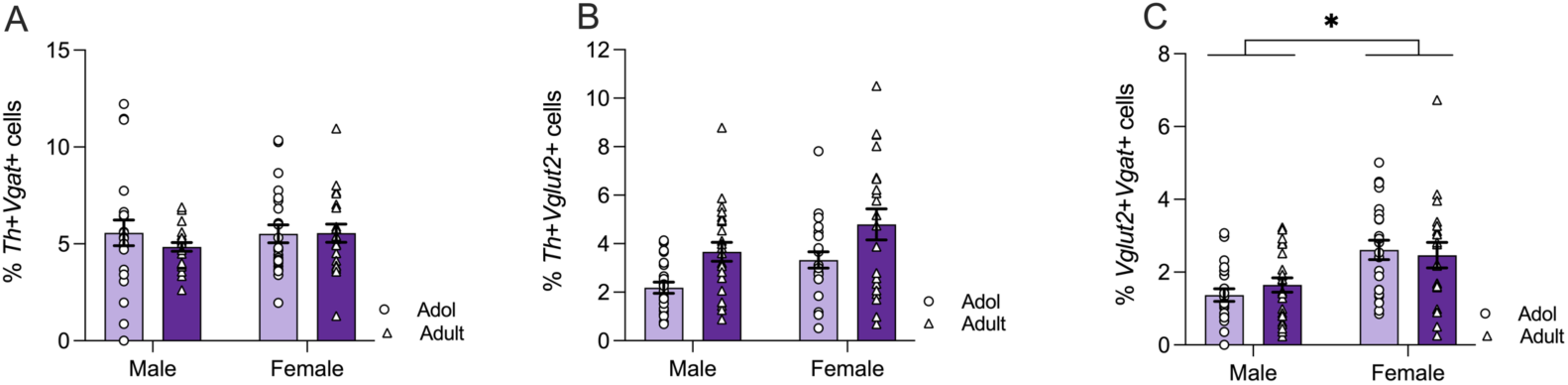
**(A)** The percentage of cells in VTA ROIs that were positive for both *Th* and *Vgat* and **(B)** *Th* and *Vglut2* transcripts. **(C)** Female mice had a higher percentage of cells positive for both *Vglut2* and *Vgat* compared with males (\**P*=0.01). Data expressed as mean ± SEM with 4-8 ROI per mouse from *n*=3 mice per sex per age.

### nAChR subunit expression per single cell type in the VTA

We then examined α4, α5, α6, β2 and β3 nAChR subunit transcript expression within *Th, Vglut2, Vgat* and *Gfap* positive cells in the VTA to identify whether different cell types have different patterns of nAChR subunit expression. In VTA *Th*+ cells, an average of 95% of cells were positive for α4, 95% were positive for α6, 97% were positive for β2 and 94% were positive for β3 across all groups (**Fig. 4A**). The α5 nAChR subunit transcript was expressed in an average of 41-46% of VTA *Th*+ cells across all groups. There were no significant differences in expression between ages within sex for α4, α5, α6, β2 or β3. When age was collapsed, there was a trend for males to have a higher percentage of β3 positive *Th+* cells compared with females (nested Student’s t-test, *t*=2.11, *P*=0.06), and males had a small but significantly higher percentage of α4 positive *Th+* cells compared with females (nested Student’s t-test, *t*=2.40, *P*=0.04). Overall, nearly all VTA *Th*+ cells express α4, α6, β2 and β3 transcripts, with approximately half of the cells expressing α5, suggesting the possibility of the α4α5α6β2β3 nAChR subtype in half of VTA dopaminergic cells.

**Fig. 4.**
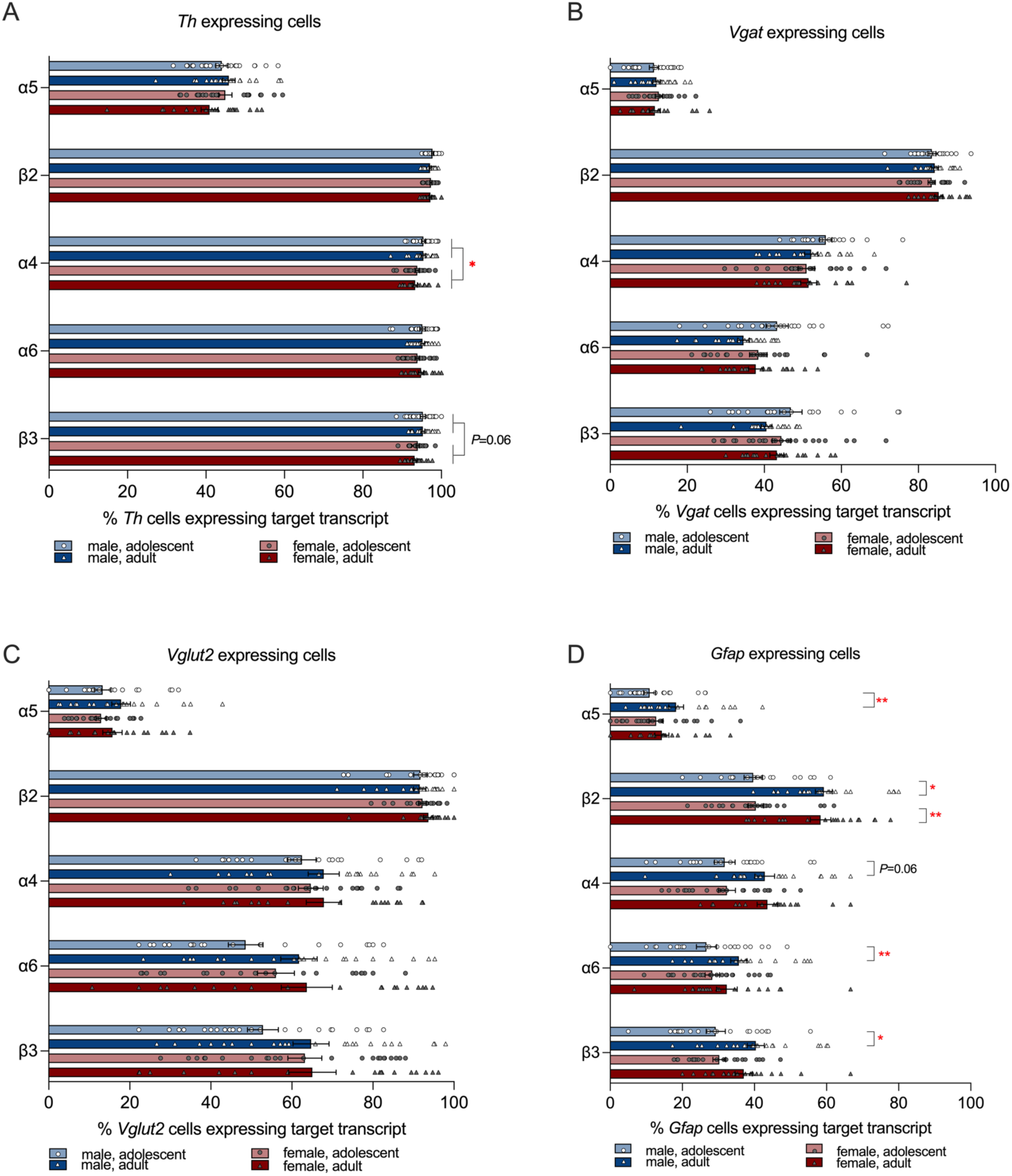
The percentage of cells in VTA ROIs expressing α4, α5, α6, β2, and β3 nAChR subunit transcripts in **(A)** *Th*+ cells, **(B)** *Vgat*+ cells, **(C)** *Vglut2*+ cells and **(D)** *Gfap*+ cells. \**P*<0.05, \*\**P*<0.01 between the indicated groups. Data expressed as mean ± SEM with 4-8 ROI per mouse from *n*=3 mice per sex per age.

We found a different expression profile in VTA *Vgat*+ cells compared with *Th*+ cells. The β2 transcript was the most prevalent subunit and was expressed in an average of 84% of *Vgat*+ cells across all groups (**Fig. 4B**). Interestingly, the percentage of *Vgat*+ cells that were positive for the α4 transcript was only 51-56%. For α6 and β3, which are often found together in the same receptor subtype, the %*Vgat+* cells that were positive was also lower compared with *Th+* cells, with 41-47% of *Vgat+* cells positive for β3 and 35-44% positive for α6. The α5 transcript was the lowest expressing subunit, with only 12-13% of *Vgat*+ cells positive for this transcript. There were no significant differences in subunit transcript expression in *Vgat*+ cells.

The nAChR subunit transcript expression profile in *Vglut2*+ cells was similar to *Vgat*+ cells, with the β2 subunit transcript expressed in an average of 92-95% of *Vglut2*+ cells across all groups (**Fig. 4C**). The α4, α6 and β3 subunit transcripts were expressed in fewer cells, with the α4 subunit transcript in 62-67%, α6 in 48-61% and β3 in 53-64% of *Vglut2+* cells. The α5 transcript had the lowest expression and was detected in 13-18% of *Vglut2*+ cells. There were no significant differences in subunit transcript expression in *Vglut2*+ cells.

We next examined *Gfap*+ cells, a marker of astrocytes, in the VTA. The β2 nAChR subunit was the most prevalent subunit, however it was expressed in only 40-59% of *Gfap*+ cells (**Fig. 4D**). There was a significant difference in expression between male adolescents and male adults, with adults having greater prevalence of β2 in astrocytes compared with adolescents (adolescent males 39.7 ± 2.6% vs adult males 59.3 ± 2.4%, nested Student’s *t*-test, *t*=4.07, *P*=0.02). An age effect was also observed for α6, β3 and α5 in male mice, with adults having greater prevalence of these subunits in astrocytes compared with adolescents (α6: adolescents 25.5 ± 2.9% vs 35.7 ± 2.3% adults, *t*=2.79, *P*=0.008; β3: adolescents 27.9 ± 2.9% vs adults 40.4 ± 2.4%, *t*=2.98, *P*=0.04; α5: adolescents 10.9 ± 1.7% vs adults 18.3 ± 2.1%, t=2.73, *P*=0.009, all nested Student’s *t*-tests). There was also a strong trend toward greater prevalence of the α4 subunit in astrocytes in adult males compared with adolescent males (adolescent males 31.8 ± 2.9% vs adult males 42.9 ± 2.7%, nested Student’s *t*-test, *t*=2.63, *P*=0.06). An age effect was also observed for β2 in female mice (adolescent females 40.4 ± 2.1% vs adult females 58.3 ± 2.8%, nested Student’s *t*-test, *t*=4.89, *P*=0.008). There was a trend for greater prevalence of α4 and β3 nAChR subunit transcripts in adult females compared with adolescents. Overall, adult mice had more prevalent astrocytic expression of nAChR subunit transcripts compared with adolescents, particularly in male mice.

### Single and co-expressing cell types in the LC

We first determined whether ROIs from the LC contained an equal number of cells across all groups. The average number of cells per LC ROI was 206.3 ± 112.3 (mean ± SD), with no differences across groups (nested one-way ANOVA, F=0.70, *P*=0.58, n=2-7 ROI per mouse, from *n*=3 mice per sex per age). We then examined cell type expression in the LC starting with noradrenergic cells. The percentage of cells per ROI expressing *Th* (%*Th*+), a marker of noradrenergic cells in the LC, was 32.6 ± 1.7% in adolescent males, 36.7 ± 2.1% in adult males, 32.3 ± 2.3% in adolescent females, and 41.5 ± 1.7% in adult females (**Fig. 5A**). The %*Vgat+* cells was 6.3 ± 1.7% in adolescent males, 6.7 ± 1.1% in adolescent females, 7.3 ± 1.4% in adult males and 6.5 ± 1.3% in adult females, **Fig. 5B**. The %*Vglut2+* cells was 8.3 ± 0.6% in adolescent males, 8.3 ± 0.9% in adolescent females, 10.4 ± 1.3% in adult males and 8.0 ± 1.1% in adult females, **Fig. 5C**. The %*Gfap*+ cells in the LC were 7.2 ± 1.9% in adolescent males, 11.8 ± 1.9% in adolescent females, 8.2 ± 2.2% in adult males and 8.3 ± 2.4% in adult females, **Fig. 5D**. We found a small percentage of LC cells that co-expressed two cell-type markers. The percentage of %*Th*+*Vgat*+ cells was 0.7 ± 0.2% in adolescent males, 1.8 ± 0.2% in adult males, 1.3 ± 0.3% in adolescent females and 2.2 ± 0.3 % in adult females (**Fig. 6A**). There was a trend for greater prevalence of %*Th+Vgat+* cells in male adults compared with male adolescents (nested Student’s *t*-test, *t*=2.353, *P*=0.08). The percentage of *Th*+*Vglut2*+ was 1.2 ± 0.3% in adolescent males, 1.4 ± 0.2% in adult males, 1.1 ± 0.2% in adolescent females and 1.7 ± 0.4% in adult females (**Fig. 6B**). Less than one percent of cells co-expressed *Vgat* and *Vglut2*, with 0.5 ± 0.2% in adolescent males, 0.8 ± 0.2% in adult males, 0.7 ± 0.2% in adolescent females and 0.5 ± 0.2% in adult females (**Fig. 6C**). Overall, there were no significant differences in the prevalence of single or co-expressing cell types examined across groups in the LC.

**Fig. 5.**
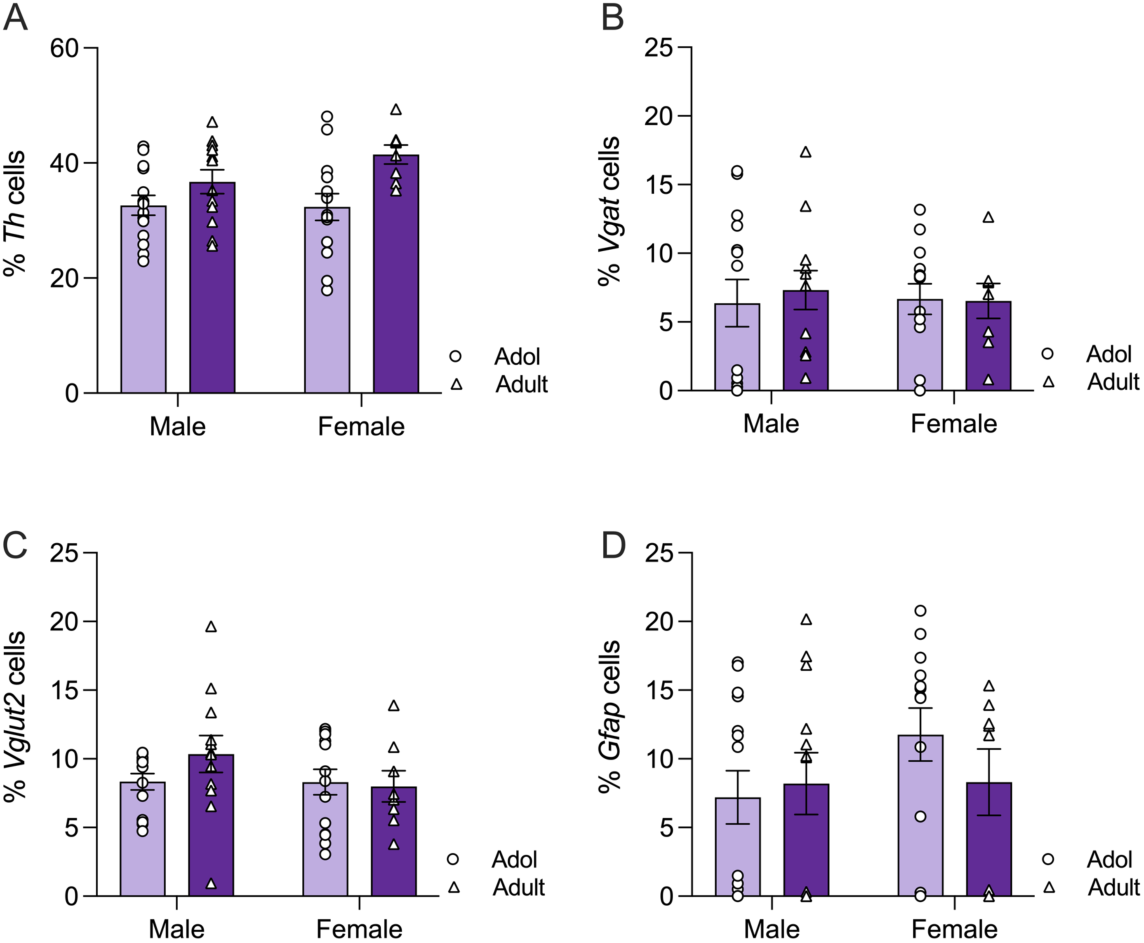
The percentage of cells in LC ROI that were positive for **(A)** *Th*, **(B)** *Vgat,* **(C)** *Vglut2* and **(D)** *Gfap* transcript. Data expressed as mean ± SEM, with 2-7 ROI per mouse from *n*=3 mice per sex per age.

**Fig. 6.**
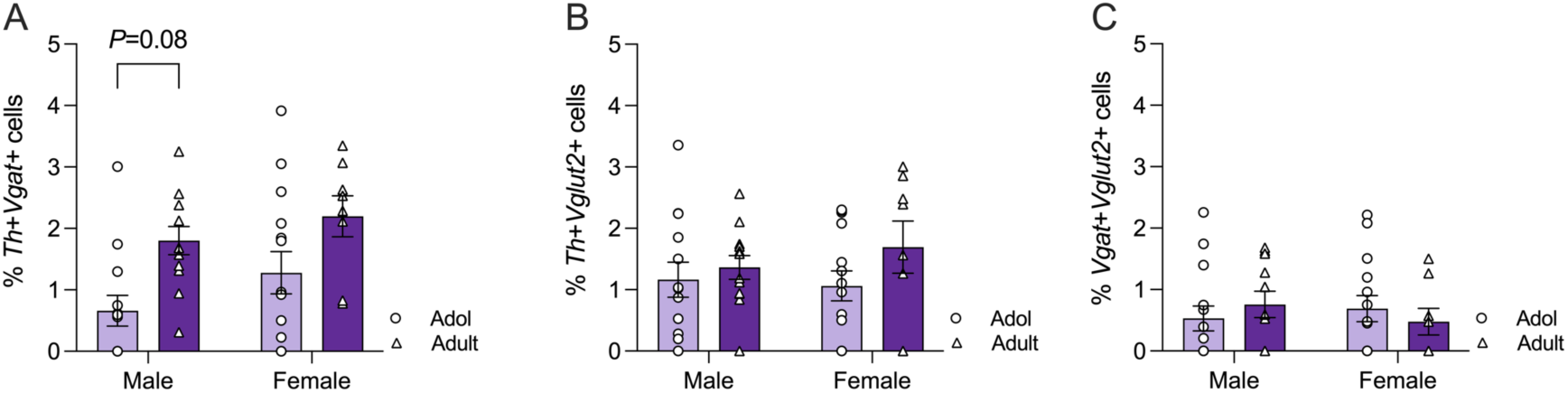
The percentage of cells in LC ROI positive for **(A)** *Th* and *Vgat*, **(B)** *Th* and *Vglut2*, and **(C)** *Vgat* and *Vglut2*. Data expressed as mean ± SEM, with 2-7 ROI per mouse from *n*=3 mice per sex per age.

### nAChR subunit expression per single cell type in the LC

We then examined nAChR subunit transcript expression profiles in the different cell types in the LC. We found a distinct profile of nAChR subunit expression in LC noradrenergic cells that suggest the presence of a α6(non-α4)α3β2 receptor subtype. In addition, the α5 subunit transcript was minimally detected in all cell types, suggesting absence of nAChRs containing this subunit in the LC. For LC noradrenergic cells identified by *Th*+, the β2 nAChR subunit transcript had the highest prevalence with an average of 91-94% prevalence. Adult males had a higher %*Th+* cells expressing β2 compared with adolescent males (adult males 94.9 ± 0.9% vs adolescent males 90.8 ± 1.0%, nested Student’s *t*-test, *t*=3.04, *P*=0.006, **Fig. 7A**). This age effect was also observed in female mice (adult females 93.9 ± 0.7% vs adolescent females 90.5 ± 0.9%, nested Student’s *t*-test, *t*=2.49, *P*=0.02). In contrast to β2, the α4 subunit transcript was expressed at low prevalence with only 7-8% of *Th+* cells positive for α4, with no significant differences between groups. The α6 and β3 subunit transcripts were far more prevalent in *Th*+ cells than α4 with an average of 88-93% for α6 and 81-89% for β3, with no significant difference in prevalence across groups. The α5 transcript was minimally detected with less than 1% of *Th*+ cells expressing this transcript in all groups.

**Fig. 7.**
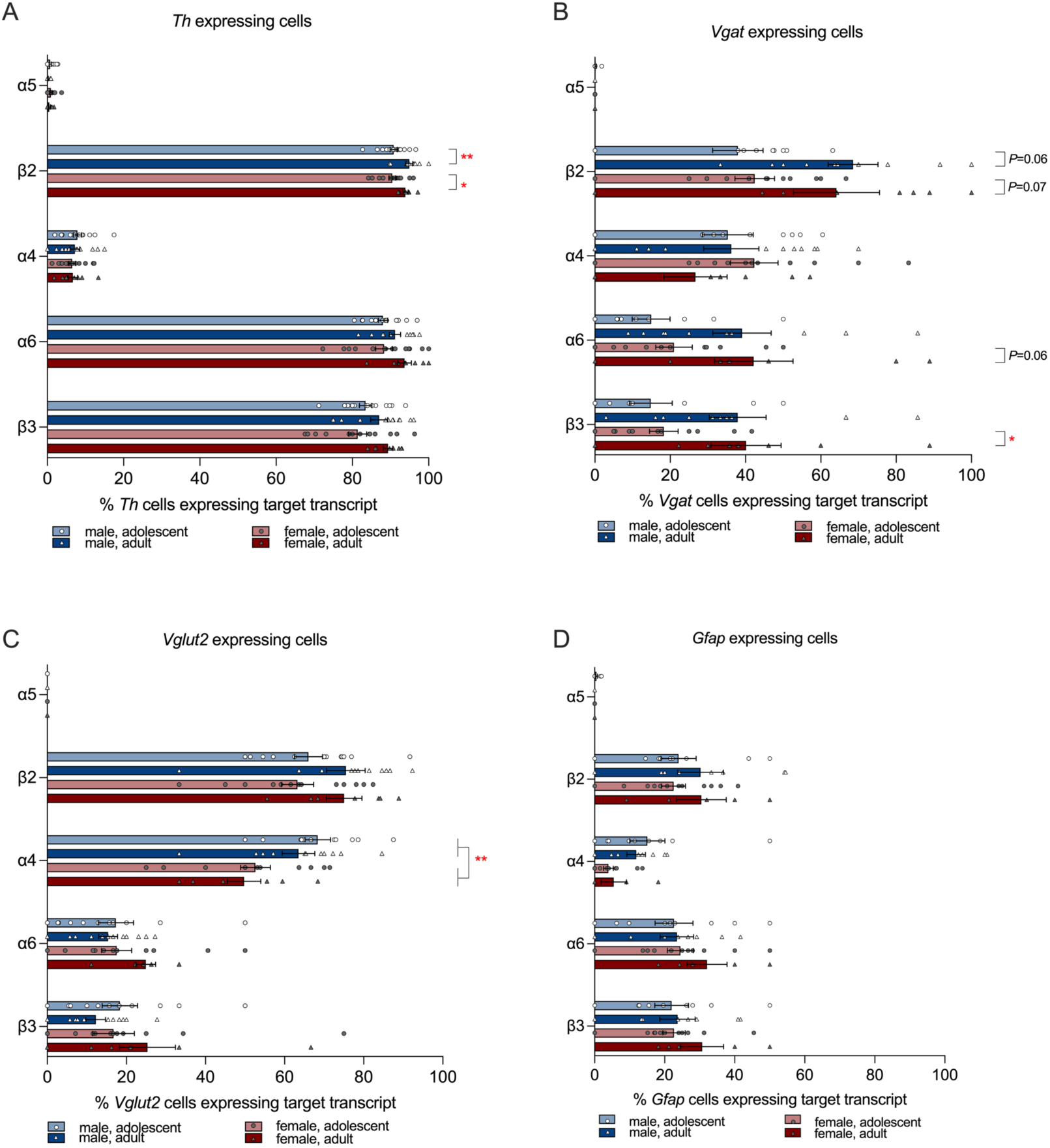
The percentage of cells in LC ROIs expressing α4, α5, α6, β2, and β3 nAChR subunit transcripts in **(A)** *Th*+ cells, **(B)** *Vgat*+ cells, **(C)** *Vglut2*+ cells and **(D)** *Gfap*+ cells. \**P*<0.05, \*\**P*<0.01 between the indicated groups. Data expressed as mean ± SEM with 2-7 ROI per mouse from *n*=3 mice per sex per age.

The expression of nAChR subunit transcripts in LC *Vgat*+ cells was more variable compared with *Th*+ cells. There was a trend for an age effect for β2 transcript prevalence in male mice, with 43.6 ± 8.3% of *Vgat+* cells positive for β2 in adolescent males and 71.2 ± 6.6% in adult males (nested Student’s *t*-test, *t*=2.53, *P*=0.06, **Fig. 7B**). There was a similar trend in females, with 42.4 ± 5.3% of *Vgat+* cells positive for β2 in adolescent females and 64.2 ± 11.4% in adult females (nested Student’s *t*-test, *t*=1.93, *P*=0.07). The α6 subunit was found in 14.9 ± 5.0% of *Vgat+* cells in adolescent males, 44.1 ± 8.7% in adult males, 21.0 ± 4.8% in adolescent females and 42.2 ± 10.4% in adult females. Prevalence of the α6 subunit transcript in *Vgat+* cells was also variable. There was a trend for female adults to have greater prevalence compared with adolescents (females: nested Student’s *t*-test, *t*=2.05, *P*=0.06; males: nested Student’s *t*-test*, t*=1.47, *P*=0.21). The β3 subunit transcript showed a similar pattern of expression compared with α6. There was a significant difference in β3 prevalence in *Vgat+* cells between female adolescents and adults (nested Student’s *t*-test*, t*=2.47, *P*=0.02) but not in males (nested Student’s *t*-test*, t*=1.49, *P*=0.21). The α4 subunit was expressed in an average of 27-42% of *Vgat+* cells, with no significant differences in prevalence across groups. The α5 subunit transcript was minimally detected with less than 0.5% of *Vgat+* cells expressing this transcript. Overall, these data suggest that nAChR transcripts may be more prevalent in LC GABA neurons in adult mice compared with adolescents, particularly in female mice.

In LC *Vglut2*+ cells, the β2 and α4 subunits were the most prevalent transcripts. An average of 63-75% of *Vglut2+* cells were positive for the β2 subunit with no significant differences in prevalence across groups. The α4 subunit present in 50-68% of *Vglut2+* cells. There was no effect of age in either sex, and when age was collapsed males had greater prevalence of α4 in *Vglut2*+ cells compared with females (nested Student’s *t*-test*, t*=3.56, *P*=0.005, **Fig. 7C**). The α6 and β3 subunit transcripts had similar levels of expression, with 15- 25% of *Vglut2*+ cells expressing α6 and 12-25% expressing β3, with no differences in prevalence across groups. The α5 subunit transcript was not detected in *Vglut2+* cells in any group. Overall, the nAChR expression profile in both LC glutamatergic and GABAergic cells suggests that the α4α6β2β3 subtype is possible.

LC astrocytes (*Gfap*+ cells) had lower prevalence of β2 compared with LC neuronal cell types with an average of 23-30% prevalence across all groups (**Fig. 7D**). The α4 subunit was expressed in only 4-12%, the α6 transcript in 23-32% and β3 in 22-31% of *Gfap+* cells. The α5 transcript was not detected in *Gfap+* cells in any group. Unlike *Gfap*+ cells in the VTA, there were no significant differences in nAChR transcript prevalence between age in either sex.

## Discussion

There were no significant differences in the number of cells per ROI across all groups for the VTA and LC, which suggests that differences in the percentage of cells that were positive for the target transcripts most likely represent changes in the number of cells expressing these transcripts. A limitation of this approach is that the cell counts for the co-expressing cell types (e.g. *Th*+*Vglut2*+) are included in the cell counts for the single cell-type markers (e.g. *Th*+) as these could not be separated.

Our VTA ROI was an identically sized rectangle that encompassed the medial VTA where we found that ∼30% cells were dopaminergic, ∼20% of cells were GABAergic and 6-9% of cells were glutamatergic. These cell type percentages agree with previous reports of dopamine, GABA and glutamate cells in medial sections of rodent VTA (Nair-Roberts et al., 2008; Yamaguchi et al., 2011; Morales and Margolis, 2017), thus providing confidence in this unbiased approach. In the VTA, we found sex effects in the prevalence of glutamatergic cells and co-expressing glutamatergic+GABAergic cells with female mice having a greater proportion of these cells in the VTA compared with males. Co-expressing glutamatergic+GABAergic cells have been previously documented in the VTA in male rodents (Root et al., 2014; Root et al., 2018b) but to our knowledge this is the first report of these co-expressing cells in female animals; moreover, these data indicate that these co-expressing cells may be more prevalent in females. As VTA glutamatergic neurons are involved in salience, reward and aversion (Wang et al., 2015; Root et al., 2018a), these data suggest that glutamatergic involvement in these responses may be more prominent in females due to the greater expression of these cells in the VTA in females compared with males.

Noradrenergic neurons and noradrenaline signaling mechanisms are implicated in attention, aging, neurodegeneration, stress responses, SUD and AUD (Poe et al., 2020; Beardmore et al., 2021; Downs and McElligott, 2022). The LC provides broad noradrenergic innervation in the brain and nAChR regulation of LC cells has not been well documented. We used *Th*, which is required for synthesis of dopamine, adrenaline and noradrenaline, as a marker of noradrenergic cells in the LC. Although the dopaminergic and noradrenergic circuits in the brain are largely dissociated, it may be possible that noradrenergic neurons in the LC can also produce dopamine (Ranjbar-Slamloo and Fazlali, 2019). We found that noradrenergic LC cells (*Th*+ cells) comprised ∼30-40% of all cells, with GABA and glutamate cells at ∼6-10% of cells. We also detected cells that co-expressed neuronal markers in the LC, suggesting that cells may synthesize both noradrenaline and glutamate, noradrenaline and GABA, or glutamate and GABA in the LC. In contrast to the VTA, where neurons co-expressing cell-type markers have been examined for their roles in neurobiology (Root et al., 2016; Morales and Margolis, 2017; Zell et al., 2020), the physiological role of these co-expressing cells in the LC have not been well studied.

In addition to neuronal cell markers, we probed for *Gfap*, a marker of astrocytes, in both the VTA and LC. Astrocytes are important regulators of synaptic function and increasing evidence suggests that astrocytes mediate aspects of SUD including drug seeking, extinction and relapse (Giacometti et al., 2020; Wang et al., 2022). We found that 7-10% of the cells were astrocytes in both brain regions. In the VTA, adult males had a higher prevalence of astrocytes compared with adolescent males, indicating a potentially greater role for astrocytes in adults. However, no age dependent differences in astrocyte prevalence were found in the LC in either sex, suggesting that developmental astrocyte expression differs by brain region.

We then identified the expression profile of five nAChR subunit transcripts, α4, α5, α6, β2 and β3, in different cell types in the VTA and LC in adult and adolescent, male and female C57BL/6J mice. Our findings highlight the variation in nAChR subunit transcript expression profiles in different cell types and brain regions, and illustrate the potential for complex and differential nAChR signaling. We did not probe for all eleven nAChR subunit transcripts as we focused on the transcripts that were heavily expressed in the VTA and LC, and are thought to form high-affinity nAChRs containing the α4 and/or α6 subunit (Whiteaker et al., 2000; Marks et al., 2006; Wills et al., 2022). Our study identifies the percentage of a cell type that is positive for a nAChR transcript and does not account for the magnitude of the target transcript expression in positive cells. In addition, the data on any one transcript is independent from the other transcripts and as such does not identify the composition of the nAChR pentamer in a cell. The composition of nAChR pentamers, also called the nAChR subtype, is more difficult to ascertain as nAChR subunits can combine into produce different receptor subtypes and even different stoichiometries of receptor subtypes. However, identifying the presence of subunit transcripts in specific cell types is useful as it provides an indication of which nAChR subtype combinations are possible (Azam et al., 2002; Yan et al., 2018). These data also further the interpretation of past nAChR subunit knock-out studies that used transgenic mice with constitutive, global deletions of the α4, β2 or α6 subunit, which have shown that deletion of these subunits lead to altered nicotine self-administration, alcohol consumption, and nicotine and alcohol reward (Picciotto et al., 1998; Pons et al., 2008; Hendrickson et al., 2011; Liu et al., 2013; Guildford et al., 2016). Identifying α4, α6 and β2 nAChR subunit expression in lesser studied neuronal cell types and astrocytes in multiple brain regions highlights the potential contribution of these cells and brain regions to cholinergic mechanisms of AUD and SUD. In addition, identifying nAChR subunit expression in specific cell types and brain regions may provide novel druggable targets for nAChR based pharmacotherapies for AUD and SUD, such as targeting α6* nAChRs in LC noradrenergic neurons and astrocytes in both the VTA and LC.

We found that over 90% of VTA *Th*+ cells were positive for the α4, α6, β2 and β3 transcripts, suggesting that nearly all VTA dopamine neurons express these four transcripts. These data replicate prior findings that α4, α6, β2 and β3 are heavily expressed in VTA dopamine neurons, which is the cell type most studied in regards to nAChR subunit expression in the VTA (Azam et al., 2002; Champtiaux et al., 2003; Azam et al., 2007; Yang et al., 2009), again providing confidence in this unbiased approach. We found that ∼50% of VTA dopaminergic cells expressed α5, suggesting that half of VTA dopamine neurons may have a nAChR composition of α4α5α6β2β3 and half with α4α6β2β3. Interestingly, all five nAChR subunits were also detected in VTA glutamatergic and GABAergic cells. Although α4 and β2 are widely expressed, the α6 and β3 subunits were previously thought to be limited to VTA dopamine neurons (Charpantier et al., 1998). We found that α6 and β3 were expressed in approximately half of all VTA glutamatergic and GABAergic cells. Although the level of α6 and β3 expression in GABA cells is much lower compared with dopamine neurons, our data adds to the recent studies that have detected and uncovered a role for α6* on GABA neurons in the regulation of alcohol effects, highlighting the involvement of nAChRs in drug effects even when expressed at low levels (Steffensen et al., 2018; Wadsworth et al., 2023; Yan et al., 2018).

Noradrenergic signaling and the LC have been implicated in mediating SUD and AUD mechanisms (Downs and McElligott, 2022). Cholinergic regulation of the LC has not been a heavy focus of SUD research and the nAChR expression profile in the LC has not been extensively studied. Past studies in adolescent rats have shown that LC cells express α3, α5, α6, β3 and β4 transcripts, however expression was not partitioned by cell type (Lena et al., 1999). We found that the nAChR subunit expression pattern in LC noradrenergic neurons was different compared with VTA dopaminergic cells. LC noradrenergic, glutamatergic and GABAergic cells expressed α4, α6, β2 and β3, but α5 was nearly undetectable in all three neuronal cell-types. Moreover, 80-90% of noradrenergic cells were positive for α6, β2, and β3 expression; however, less than 10% were positive for α4. This suggests that non-α4 nAChR subtypes, such as the α6(non-α4)β2β3 subtype, may be the predominant receptor in LC noradrenergic cells. In contrast, LC GABAergic and glutamatergic neurons may express both the α4β2 and α4α6β2β3 nAChR subtypes as the prevalence of α4 was much higher in both cell types. Overall, we identified possible nAChR subtypes in LC noradrenergic, GABAergic and glutamatergic cells, and found distinct differences in nAChR subunit transcript expression profiles between the VTA and LC.

Astrocytes are important regulators of synaptic function and support neurotransmitter homeostasis. Astrocytes encompass the pre- and post-synaptic neuronal terminals forming a complex referred to as the tripartite synapse. Recent evidence suggests that astrocytes may have a more active role in the neurobiology of SUD than previously considered (Giacometti et al., 2020; Wang et al., 2022). Astrocytes express neurotransmitter receptors and past studies have identified the α4, α7, β2 and β3 nAChR subunit transcripts in rat astrocyte cultures (Xiu et al., 2005). We find that mouse astrocytes expressed α4, α6, β2 and β3 nAChR subunit transcripts, suggesting that astrocytes may express multiple subtypes including the high affinity α4* and/or α6* nAChR subtype, potentially leading to complex cholinergic regulation of astrocyte function. Intriguingly, the expression profile differed between astrocytes in the VTA compared with the LC, as the α5 subunit was present in VTA astrocytes but undetectable in LC astrocytes, suggesting that nAChR subtype expression on VTA astrocytes are likely distinct from LC astrocytes. Moreover, we found significant effects of age in the expression of nAChR transcripts in VTA astrocytes but not in LC astrocytes. Male and female adult mice had a significantly higher prevalence of nAChR subunit expression in VTA astrocytes compared with adolescent mice. Age related changes in nAChR expression in the VTA may be particularly relevant for SUD as adolescence is a period of heightened vulnerability to developing SUD, AUD and nicotine use (Spear, 2000; Laviolette, 2021).

Prior studies have examined age-related differences in nAChR expression in VTA neurons, but not astrocytes, and these studies have found mixed effects. One study assessed fluorescently tagged α4 subunit protein expression and observed greater fluorescence intensity at 54 days compared with 45 days in dopaminergic and GABAergic cells in the VTA in male mice (Renda et al., 2016). Our data captures transcript positivity and not the amount or intensity of expression within cells, thus changes in amount in already positive cells are not measured. Another study examined nAChR subunit transcript expression in putative VTA and substantia nigra dopamine cells in rats and found greater α6 transcript levels, with no differences in α4, β2 and β3, in adolescent rats compared with adults (Azam et al., 2007).

In summary, we found complex patterns of nAChR subunit transcript expression and prevalence in C57BL/6J mice. Prevalence of nAChR subunits differed between cell types within the same brain region, and between the same cell type across the VTA and LC. These expression patterns enable us to infer the nAChR subtype combinations that are possible in specific cell types, such as the α6(non-α4)β2β3 subtype in LC noradrenergic cells. These differences in nAChR expression across cell type and brain region may be important targets in the design of nAChR pharmacotherapies for SUD and other conditions.

## Author contribution statement

Runbo Gao wrote the MATLAB program, collected the VTA and LC data, analyzed the data, and wrote part of the methods. Amelia Schneider collected LC data and edited the manuscript. Sarah Mulloy wrote and edited the manuscript. Anna Lee conceptualized the project, analyzed the data, wrote and edited the manuscript.

## Abbreviations

nAChR: nicotinic acetylcholine receptor;
VTA: ventral tegmental area;
LC: locus coeruleus;
SUD: substance use disorder;
AUD: alcohol use disorder;
ROI: region of interest;
SEM: standard error of the mean;
SD: standard deviation;
DAPI: 4’,6-diamidino-2-phenylindole;
Vglut2: vesicular glutamate transporter 2;
Vgat: vesicular GABA transporter;
Th: tyrosine hydroxylase;
Gfap: glial fibrillary acidic protein.

## Acknowledgements and Funding

We thank Dr. Ying Zhang (Informatics Analyst, Minnesota Supercomputing Institute, University of Minnesota) for her assistance with statistical analysis. This work was supported by NIH R01 AA026598 (AML).

## References

Albuquerque, E.X., Pereira, E.F., Alkondon, M., Rogers, S.W., 2009. Mammalian nicotinic acetylcholine receptors: from structure to function. Physiol Rev 89, 73–120. http://pubmed.ncbi.nlm.nih.gov/19126755.

Azam, L., Chen, Y., Leslie, F.M., 2007. Developmental regulation of nicotinic acetylcholine receptors within midbrain dopamine neurons. Neuroscience 144, 1347–1360. http://pubmed.ncbi.nlm.nih.gov/17197101.

Azam, L., Winzer-Serhan, U.H., Chen, Y., Leslie, F.M., 2002. Expression of neuronal nicotinic acetylcholine receptor subunit mRNAs within midbrain dopamine neurons. J Comp Neurol 444, 260–274. http://pubmed.ncbi.nlm.nih.gov/11840479.

Beardmore, R., Hou, R., Darekar, A., Holmes, C., Boche, D., 2021. The Locus Coeruleus in Aging and Alzheimer’s Disease: A Postmortem and Brain Imaging Review. J Alzheimers Dis 83, 5–22. https://pubmed.ncbi.nlm.nih.gov/34219717.

Bordia, T., Hrachova, M., Chin, M., McIntosh, J.M., Quik, M., 2012. Varenicline Is a Potent Partial Agonist at alpha6beta2* Nicotinic Acetylcholine Receptors in Rat and Monkey Striatum. J Pharmacol Exp Ther 342, 327–334. http://www.ncbi.nlm.nih.gov/pubmed/22550286.

Carpenter, A.E., Jones, T.R., Lamprecht, M.R., Clarke, C., Kang, I.H., Friman, O., Guertin, D.A., Chang, J.H., Lindquist, R.A., Moffat, J., Golland, P., Sabatini, D.M., 2006. CellProfiler: image analysis software for identifying and quantifying cell phenotypes. Genome Biol 7, R100. https://pubmed.ncbi.nlm.nih.gov/17076895.

Champtiaux, N., Gotti, C., Cordero-Erausquin, M., David, D.J., Przybylski, C., Lena, C., Clementi, F., Moretti, M., Rossi, F.M., Le Novere, N., McIntosh, J.M., Gardier, A.M., Changeux, J.P., 2003. Subunit composition of functional nicotinic receptors in dopaminergic neurons investigated with knock-out mice. J Neurosci 23, 7820–7829. http://pubmed.ncbi.nlm.nih.gov/12944511.

Champtiaux, N., Han, Z.Y., Bessis, A., Rossi, F.M., Zoli, M., Marubio, L., McIntosh, J.M., Changeux, J.P., 2002. Distribution and pharmacology of alpha 6-containing nicotinic acetylcholine receptors analyzed with mutant mice. J Neurosci 22, 1208–1217. http://pubmed.ncbi.nlm.nih.gov/11850448.

Charpantier, E., Barneoud, P., Moser, P., Besnard, F., Sgard, F., 1998. Nicotinic acetylcholine subunit mRNA expression in dopaminergic neurons of the rat substantia nigra and ventral tegmental area. Neuroreport 9, 3097–3101. http://pubmed.ncbi.nlm.nih.gov/9804323.

De Biasi, M., Dani, J.A., 2011. Reward, addiction, withdrawal to nicotine. Annu Rev Neurosci 34, 105–130. http://pubmed.ncbi.nlm.nih.gov/21438686.

Downs, A.M., McElligott, Z.A., 2022. Noradrenergic circuits and signaling in substance use disorders. Neuropharmacology 208, 108997. https://pubmed.ncbi.nlm.nih.gov/35176286.

Galvin, V.C., Arnsten, A.F.T., Wang, M., 2020. Involvement of Nicotinic Receptors in Working Memory Function. Curr Top Behav Neurosci 45, 89–99. https://pubmed.ncbi.nlm.nih.gov/32451954.

Giacometti, L.L., Chandran, K., Figueroa, L.A., Barker, J.M., 2020. Astrocyte modulation of extinction impairments in ethanol-dependent female mice. Neuropharmacology 179, 108272. https://pubmed.ncbi.nlm.nih.gov/32801026.

Gotti, C., Guiducci, S., Tedesco, V., Corbioli, S., Zanetti, L., Moretti, M., Zanardi, A., Rimondini, R., Mugnaini, M., Clementi, F., Chiamulera, C., Zoli, M., 2010. Nicotinic acetylcholine receptors in the mesolimbic pathway: primary role of ventral tegmental area alpha6beta2* receptors in mediating systemic nicotine effects on dopamine release, locomotion, and reinforcement. J Neurosci 30, 5311–5325. http://pubmed.ncbi.nlm.nih.gov/20392953.

Grady, S.R., Salminen, O., Laverty, D.C., Whiteaker, P., McIntosh, J.M., Collins, A.C., Marks, M.J., 2007. The subtypes of nicotinic acetylcholine receptors on dopaminergic terminals of mouse striatum. Biochem Pharmacol 74, 1235–1246. http://pubmed.ncbi.nlm.nih.gov/17825262.

Guildford, M.J., Sacino, A.V., Tapper, A.R., 2016. Modulation of ethanol reward sensitivity by nicotinic acetylcholine receptors containing the α6 subunit. Alcohol 57, 65–70. https://pubmed.ncbi.nlm.nih.gov/27793544.

Hendrickson, L.M., Gardner, P., Tapper, A.R., 2011. Nicotinic acetylcholine receptors containing the alpha4 subunit are critical for the nicotine-induced reduction of acute voluntary ethanol consumption. Channels (Austin) 5, 124–127. http://pubmed.ncbi.nlm.nih.gov/21239887.

Hendrickson, L.M., Guildford, M.J., Tapper, A.R., 2013. Neuronal nicotinic acetylcholine receptors: common molecular substrates of nicotine and alcohol dependence. Front Psychiatry 4, 1–16. http://pubmed.ncbi.nlm.nih.gov/23641218.

Hurst, R., Rollema, H., Bertrand, D., 2013. Nicotinic acetylcholine receptors: from basic science to therapeutics. Pharmacol Ther 137, 22–54. http://pubmed.ncbi.nlm.nih.gov/22925690.

Hynes, T.J., Hrelja, K.M., Hathaway, B.A., Hounjet, C.D., Chernoff, C.S., Ebsary, S.A., Betts, G.D., Russell, B., Ma, L., Kaur, S., Winstanley, C.A., 2021. Dopamine neurons gate the intersection of cocaine use, decision making, and impulsivity. Addict Biol 26, e13022. https://pubmed.ncbi.nlm.nih.gov/33559379.

Kamentsky, L., Jones, T.R., Fraser, A., Bray, M.A., Logan, D.J., Madden, K.L., Ljosa, V., Rueden, C., Eliceiri, K.W., Carpenter, A.E., 2011. Improved structure, function and compatibility for CellProfiler: modular high-throughput image analysis software. Bioinformatics 27, 1179–1180. https://pubmed.ncbi.nlm.nih.gov/21349861.

Laviolette, S.R., 2021. Molecular and neuronal mechanisms underlying the effects of adolescent nicotine exposure on anxiety and mood disorders. Neuropharmacology 184, 108411. https://pubmed.ncbi.nlm.nih.gov/33245960.

Le Novere, N., Zoli, M., Changeux, J.P., 1996. Neuronal nicotinic receptor alpha 6 subunit mRNA is selectively concentrated in catecholaminergic nuclei of the rat brain. Eur J Neurosci 8, 2428–2439. http://pubmed.ncbi.nlm.nih.gov/8950106.

Lena, C., de Kerchove, D.A., Cordero-Erausquin, M., Le Novere, N., del Mar Arroyo-Jimenez, M., Changeux, J.P., 1999. Diversity and distribution of nicotinic acetylcholine receptors in the locus ceruleus neurons. Proc Natl Acad Sci U S A 96, 12126–12131. http://pubmed.ncbi.nlm.nih.gov/10518587.

Liu, L., Hendrickson, L.M., Guildford, M.J., Zhao-Shea, R., Gardner, P.D., Tapper, A.R., 2013. Nicotinic Acetylcholine Receptors Containing the alpha4 Subunit Modulate Alcohol Reward. Biol Psychiatry 73, 738–746. http://pubmed.ncbi.nlm.nih.gov/23141806.

Mackey, E.D., Engle, S.E., Kim, M.R., O’Neill, H.C., Wageman, C.R., Patzlaff, N.E., Wang, Y., Grady, S.R., McIntosh, J.M., Marks, M.J., Lester, H.A., Drenan, R.M., 2012. alpha6* Nicotinic Acetylcholine Receptor Expression and Function in a Visual Salience Circuit. J Neurosci 32, 10226–10237. http://pubmed.ncbi.nlm.nih.gov/22836257.

Marks, M.J., Whiteaker, P., Collins, A.C., 2006. Deletion of the alpha7, beta2, or beta4 nicotinic receptor subunit genes identifies highly expressed subtypes with relatively low affinity for [3H]epibatidine. Mol Pharmacol 70, 947–959. http://pubmed.ncbi.nlm.nih.gov/16728647.

McQuin, C., Goodman, A., Chernyshev, V., Kamentsky, L., Cimini, B.A., Karhohs, K.W., Doan, M., Ding, L., Rafelski, S.M., Thirstrup, D., Wiegraebe, W., Singh, S., Becker, T., Caicedo, J.C., Carpenter, A.E., 2018. CellProfiler 3.0: Next-generation image processing for biology. PLoS Biol 16, e2005970. https://pubmed.ncbi.nlm.nih.gov/29969450.

Mihalak, K.B., Carroll, F.I., Luetje, C.W., 2006. Varenicline is a partial agonist at alpha4beta2 and a full agonist at alpha7 neuronal nicotinic receptors. Mol Pharmacol 70, 801–805. http://www.ncbi.nlm.nih.gov/pubmed/16766716.

Morales, M., Margolis, E.B., 2017. Ventral tegmental area: cellular heterogeneity, connectivity and behaviour. Nat Rev Neurosci 18, 73–85. https://pubmed.ncbi.nlm.nih.gov/28053327.

Morel, C., Montgomery, S., Han, M.H., 2019. Nicotine and alcohol: the role of midbrain dopaminergic neurons in drug reinforcement. Eur J Neurosci 50, 2180–2200. https://pubmed.ncbi.nlm.nih.gov/30251377.

Nair-Roberts, R.G., Chatelain-Badie, S.D., Benson, E., White-Cooper, H., Bolam, J.P., Ungless, M.A., 2008. Stereological estimates of dopaminergic, GABAergic and glutamatergic neurons in the ventral tegmental area, substantia nigra and retrorubral field in the rat. Neuroscience 152, 1024–1031. https://pubmed.ncbi.nlm.nih.gov/18355970.

Nestler, E.J., 2005. Is there a common molecular pathway for addiction? Nat Neurosci 8, 1445– 1449. http://pubmed.ncbi.nlm.nih.gov/16251986.

Ngolab, J., Liu, L., Zhao-Shea, R., Gao, G., Gardner, P.D., Tapper, A.R., 2015. Functional Upregulation of α4* Nicotinic Acetylcholine Receptors in VTA GABAergic Neurons Increases Sensitivity to Nicotine Reward. J Neurosci 35, 8570–8578. http://pubmed.ncbi.nlm.nih.gov/26041923.

Picciotto, M.R., Addy, N.A., Mineur, Y.S., Brunzell, D.H., 2008. It is not “either/or”: activation and desensitization of nicotinic acetylcholine receptors both contribute to behaviors related to nicotine addiction and mood. Prog Neurobiol 84, 329–342. http://pubmed.ncbi.nlm.nih.gov/18242816.

Picciotto, M.R., Zoli, M., Rimondini, R., Lena, C., Marubio, L.M., Pich, E.M., Fuxe, K., Changeux, J.P., 1998. Acetylcholine receptors containing the beta2 subunit are involved in the reinforcing properties of nicotine. Nature 391, 173–177. http://pubmed.ncbi.nlm.nih.gov/9428762.

Poe, G.R., Foote, S., Eschenko, O., Johansen, J.P., Bouret, S., Aston-Jones, G., Harley, C.W., Manahan-Vaughan, D., Weinshenker, D., Valentino, R., Berridge, C., Chandler, D.J., Waterhouse, B., Sara, S.J., 2020. Locus coeruleus: a new look at the blue spot. Nat Rev Neurosci 21, 644–659. https://pubmed.ncbi.nlm.nih.gov/32943779.

Pons, S., Fattore, L., Cossu, G., Tolu, S., Porcu, E., McIntosh, J.M., Changeux, J.P., Maskos, U., Fratta, W., 2008. Crucial role of alpha4 and alpha6 nicotinic acetylcholine receptor subunits from ventral tegmental area in systemic nicotine self-administration. J Neurosci 28, 12318–12327. http://pubmed.ncbi.nlm.nih.gov/19020025.

Ranjbar-Slamloo, Y., Fazlali, Z., 2019. Dopamine and Noradrenaline in the Brain; Overlapping or Dissociate Functions. Front Mol Neurosci 12, 334. https://pubmed.ncbi.nlm.nih.gov/32038164.

Renda, A., Penty, N., Komal, P., Nashmi, R., 2016. Vulnerability to nicotine self-administration in adolescent mice correlates with age-specific expression of α4* nicotinic receptors. Neuropharmacology 108, 49–59. http://pubmed.ncbi.nlm.nih.gov/27102349.

Rollema, H., Chambers, L.K., Coe, J.W., Glowa, J., Hurst, R.S., Lebel, L.A., Lu, Y., Mansbach, R.S., Mather, R.J., Rovetti, C.C., Sands, S.B., Schaeffer, E., Schulz, D.W., Tingley, F.D., Williams, K.E., 2007. Pharmacological profile of the alpha4beta2 nicotinic acetylcholine receptor partial agonist varenicline, an effective smoking cessation aid. Neuropharmacology 52:985–994. https://pubmed.ncbi.nlm.nih.gov/17157884/

Root, D.H., Estrin, D.J., Morales, M., 2018a. Aversion or Salience Signaling by Ventral Tegmental Area Glutamate Neurons. iScience 2, 51–62. https://pubmed.ncbi.nlm.nih.gov/29888759.

Root, D.H., Mejias-Aponte, C.A., Zhang, S., Wang, H.L., Hoffman, A.F., Lupica, C.R., Morales, M., 2014. Single rodent mesohabenular axons release glutamate and GABA. Nat Neurosci 17, 1543–1551. https://pubmed.ncbi.nlm.nih.gov/25242304.

Root, D.H., Wang, H.L., Liu, B., Barker, D.J., Mód, L., Szocsics, P., Silva, A.C., Maglóczky, Z., Morales, M., 2016. Glutamate neurons are intermixed with midbrain dopamine neurons in nonhuman primates and humans. Sci Rep 6, 30615. https://pubmed.ncbi.nlm.nih.gov/27477243.

Root, D.H., Zhang, S., Barker, D.J., Miranda-Barrientos, J., Liu, B., Wang, H.L., Morales, M., 2018b. Selective Brain Distribution and Distinctive Synaptic Architecture of Dual Glutamatergic-GABAergic Neurons. Cell Rep 23, 3465–3479. https://pubmed.ncbi.nlm.nih.gov/29924991.

Spear, L., 2000. Modeling adolescent development and alcohol use in animals. Alcohol Res Health 24, 115–123. https://pubmed.ncbi.nlm.nih.gov/11199278.

Steffensen, S.C., Shin, S.I., Nelson, A.C., Pistorius, S.S., Williams, S.B., Woodward, T.J., Park, H.J., Friend, L., Gao, M., Gao, F., Taylor, D.H., Foster Olive, M., Edwards, J.G., Sudweeks, S.N., Buhlman, L.M., Michael McIntosh, J., Wu, J., 2018. α6 subunit-containing nicotinic receptors mediate low-dose ethanol effects on ventral tegmental area neurons and ethanol reward. Addict Biol 23, 1079–1093. https://pubmed.ncbi.nlm.nih.gov/pubmed/28901722.

Stirling, D.R., Swain-Bowden, M.J., Lucas, A.M., Carpenter, A.E., Cimini, B.A., Goodman, A., 2021. CellProfiler 4: improvements in speed, utility and usability. BMC Bioinformatics 22, 433. https://pubmed.ncbi.nlm.nih.gov/34507520.

Trudeau, L.E., Hnasko, T.S., Wallén-Mackenzie, A., Morales, M., Rayport, S., Sulzer, D., 2014. The multilingual nature of dopamine neurons. Prog Brain Res 211, 141–164. https://pubmed.ncbi.nlm.nih.gov/24968779.

Wadsworth, H.A., Anderson, E.Q., Williams, B.M., Ronström, J.W., Moen, J.K., Lee, A.M., McIntosh, J.M., Wu, J., Yorgason, J.T., Steffensen, S.C., 2023. Role of α6-Nicotinic Receptors in Alcohol-Induced GABAergic Synaptic Transmission and Plasticity to Cholinergic Interneurons in the Nucleus Accumbens. Mol Neurobiol https://pubmed.ncbi.nlm.nih.gov/36802012.

Wang, H.L., Qi, J., Zhang, S., Wang, H., Morales, M., 2015. Rewarding Effects of Optical Stimulation of Ventral Tegmental Area Glutamatergic Neurons. J Neurosci 35, 15948– 15954. https://pubmed.ncbi.nlm.nih.gov/26631475.

Wang, J., Holt, L.M., Huang, H.H., Sesack, S.R., Nestler, E.J., Dong, Y., 2022. Astrocytes in cocaine addiction and beyond. Mol Psychiatry 27, 652–668. https://pubmed.ncbi.nlm.nih.gov/33837268.

Whiteaker, P., Marks, M.J., Grady, S.R., Lu, Y., Picciotto, M.R., Changeux, J.P., Collins, A.C., 2000. Pharmacological and null mutation approaches reveal nicotinic receptor diversity. Eur J Pharmacol 393, 123–135. http://pubmed.ncbi.nlm.nih.gov/10771005.

Wills, L., Ables, J.L., Braunscheidel, K.M., Caligiuri, S.P.B., Elayouby, K.S., Fillinger, C., Ishikawa, M., Moen, J.K., Kenny, P.J., 2022. Neurobiological Mechanisms of Nicotine Reward and Aversion. Pharmacol Rev 74, 271–310. https://pubmed.ncbi.nlm.nih.gov/35017179.

Xiu, J., Nordberg, A., Zhang, J.T., Guan, Z.Z., 2005. Expression of nicotinic receptors on primary cultures of rat astrocytes and up-regulation of the alpha7, alpha4 and beta2 subunits in response to nanomolar concentrations of the beta-amyloid peptide(1-42). Neurochem Int 47, 281–290. https://pubmed.ncbi.nlm.nih.gov/15955596.

Yamaguchi, T., Wang, H.L., Li, X., Ng, T.H., Morales, M., 2011. Mesocorticolimbic glutamatergic pathway. J Neurosci 31, 8476–8490. https://pubmed.ncbi.nlm.nih.gov/21653852.

Yan, Y., Peng, C., Arvin, M.C., Jin, X.T., Kim, V.J., Ramsey, M.D., Wang, Y., Banala, S., Wokosin, D.L., McIntosh, J.M., Lavis, L.D., Drenan, R.M., 2018. Nicotinic Cholinergic Receptors in VTA Glutamate Neurons Modulate Excitatory Transmission. Cell Rep 23, 2236–2244. https://pubmed.ncbi.nlm.nih.gov/pubmed/29791835.

Yang, K., Buhlman, L., Khan, G.M., Nichols, R.A., Jin, G., McIntosh, J.M., Whiteaker, P., Lukas, R.J., Wu, J., 2011. Functional Nicotinic Acetylcholine Receptors Containing Alpha6 Subunits Are on GABAergic Neuronal Boutons Adherent to Ventral Tegmental Area Dopamine Neurons. J Neurosci 31, 2537–2548. http://pubmed.ncbi.nlm.nih.gov/21325521.

Yang, K., Hu, J., Lucero, L., Liu, Q., Zheng, C., Zhen, X., Jin, G., Lukas, R.J., Wu, J., 2009. Distinctive nicotinic acetylcholine receptor functional phenotypes of rat ventral tegmental area dopaminergic neurons. J Physiol 587, 345–361. http://pubmed.ncbi.nlm.nih.gov/19047205.

Zell, V., Steinkellner, T., Hollon, N.G., Warlow, S.M., Souter, E., Faget, L., Hunker, A.C., Jin, X., Zweifel, L.S., Hnasko, T.S., 2020. VTA Glutamate Neuron Activity Drives Positive Reinforcement Absent Dopamine Co-release. Neuron 107, 864–873.e4. https://pubmed.ncbi.nlm.nih.gov/32610039.

